# Measurement of atomic scattering factors by cryo-electron microscopy

**DOI:** 10.1101/2025.10.24.683059

**Authors:** Alexander Shtyrov, Hugh Wilson, Daria Slowik, Keitaro Yamashita, Jade Li, Marcin Wojdyr, Shaoxia Chen, Greg McMullan, Jude Short, Christopher J. Russo, Richard Henderson, Garib N. Murshudov

**Author notes:** GNM, RH and CJR designed research. AS, HW, DS, JL, SC, GM, JS, CJR, RH and GNM performed research. DS and RH contributed new reagents/analytic tools. KY and MW contributed code for atomic model refinement. AS, HW, JL, GM, JS, CJR, RH and GNM analysed data. AS, HW, RH and GNM wrote the paper. The authors declare no competing interests.

## Abstract

Determination of specimen structure from cryo-electron microscopy (cryo-EM) experiments relies on an accurate model of the electrostatic potential of the specimen. For biological macromolecules, the potential is strongly influenced by the presence of chemical bonds between atoms, a fact unaccounted for by models of electron scattering that are currently standard in the field. We propose a Bayesian approach to the estimation of atomic scattering factors which incorporates the effect of the molecular environment while remaining fast, interpretable and transferable between molecules. Our algorithm infers atomic scattering factors directly from maps of the electrostatic potential determined by cryo-EM single particle analysis, bypassing the need for computationally-intensive theoretical calculations. The algorithm is used to infer empirical scattering factors from high-resolution reconstructions of catalase enzymes. To illustrate its broad applicability, the algorithm is also applied to a training set of publicly-available cryo-EM data. The empirical scattering factors show improved agreement with a test set of cryo-EM reconstructions. The predictions are further validated by comparison with magnetic susceptibility values of organic compounds, as well as by application to the refinement of atomic models.

**Significance Statement:** Understanding the structure of biomolecules is key to explaining their function. Cryo-electron microscopy is a method for reconstructing the electrostatic potential distribution of a biological macro-molecule, a quantity which contains information about atomic positions and the redistribution of charge due to chemical bonding. These factors should be modelled when inferring the structure of the molecule from its electrostatic potential. We develop an improved model of the potential that takes into account chemical bonding while remaining computationally tractable. The parameters of the model are inferred from a selection of cryo-electron microscopy datasets using Bayesian methods.

---

In electron microscopy (EM), an incident beam of high-energy electrons is scattered by the electrostatic potential (ESP) of the specimen, forming a wave on its exit face (1, 2). The apparatus of an electron microscope, including the lens system, detector and subsequent digital image processing steps, is designed to recover the intensity of the exit wave as precisely as possible. Indeed, improvements in this direction, for example in electron sources, detectors and data processing algorithms, have led to significant advances in the resolution obtainable by EM (3, 4). However, the models describing the formation of the exit wave on scattering of the incident beam have received less attention compared to advances in imaging. The problem is arguably as important as image acquisition and processing, since making inferences about specimen structure requires a scattering model that is both accurate and fast to evaluate.

The present work focuses on modelling scattering from biological macromolecules. A successful scattering model has two components. The first is a procedure for generating the scattering potential of the object being imaged. The second is an approximation for calculating the scattering amplitude of the object from this potential. In EM with high-energy electrons and thin organic specimens, the second component is provided by the weak phase approximation, which states that the scattering amplitude is the Fourier transform of the scattering potential (1). However, the calculation of the scattering potential for a macromolecule is not a trivial task. For electrons, this potential is the ESP, a quantity whose precise evaluation requires calculating the electron density, then evaluating a Coulomb integral (5). Calculation of the ESP in this way is not feasible for macromolecules with current computing hardware, even less so for iterative algorithms that require multiple evaluations of the ESP and its derivatives. Therefore, when interpreting data from EM and electron diffraction (ED) experiments, it is usual to approximate the molecular ESP by a superposition of the ESPs of the individual atoms making up the molecule (2). The atoms in the molecule are in turn assumed to be identical to unbound neutral atoms in a vacuum. This series of approximations is the independent atom model (IAM). The IAM is a highly simplistic model that does not account for bonding or the presence of charged species, but has the advantage of being very fast to evaluate. Furthermore, the IAM is transferable, since the Fourier transform of the ESP can be pre-evaluated for each element. This quantity is known as the atomic scattering factor, and its value for each element is given in the International Tables for Crystallography (6). Scattering factors calculated in this way will therefore be referred to as ‘tabulated scattering factors’.

The shortcomings of the IAM as a model for electron scattering have been known for several decades, in part due to studies comparing the IAM to the results of gas-phase ED experiments. Such studies have found that the IAM matches poorly with time-averaged (7) and, more recently, time-resolved (8) measurements of electron scattering from molecules at low spatial frequency. Several attempts have been made to improve the IAM, but none has found widespread adoption. One approach is to model atoms within molecular fragments, which express the chemical intuition that a particular functional group behaves similarly in different chemical species. The resulting scattering factors are aspherical. An early example of the approach is described in (9). The transferable aspherical atom model (TAAM), based on fitting a multipole expansion into electron density maps of small molecules, is a more recent variation on this idea (10–16). The disadvantage of scattering factors based on a multipole expansion is the absence of a simple expression for the contribution of atomic displacement parameters (ADPs) to the ESP. In this case, direct summation of atomic contributions in Fourier space must be used for calculation of the ESP and its derivatives. Direct summation is significantly slower than the fast Fourier transform (FFT)- based algorithms that may be used to evaluate the ESP under the IAM (17). Some authors have instead proposed fitting spherical scattering factors in a suitable parametric form, either to experimentally-derived ESP maps (18) or to an ESP calculated computationally (19).

A different line of work aims to model the redistribution of valence electrons during bonding by assigning partial charges to selected atoms in the structure (developed in (20–23), but see also (24) and references therein). Partial charges may also be assigned to all atoms in a molecule, provided the molecule is small and the data quality is sufficiently high, as shown in (25) using ED. A drawback of such methods is that it is unclear how atomic partial charges should be represented. The studies cited above use either a linear combination of the scattering factors of the neutral atom and its ion, or point charges at the atomic nuclei, neither of which accurately reflects the redistribution of charge in the molecule.

The present contribution develops a data-driven method for the estimation of atomic scattering factors from experimentally-derived maps of the ESP distribution in macromolecules. Such maps may be derived using various techniques, but the focus here will be on reconstructions obtained by cryo-EM single particle analysis (SPA). Cryo-EM SPA is a method for determining the ESP distribution of a macromolecule in solution with a resolution in some cases high enough to see individual hydrogen atoms (26, 27).

The focus of the work is on developing computational tools for inferring scattering models from data. The algorithm outlined below is based on formulating the IAM, which is a forward model for the ESP, as an expansion over a set of basis functions. Inference of scattering factors from the ESP (the inverse problem), is then addressed within the framework of Gaussian process (GP) regression, a Bayesian nonparametric machine learning method based on specifying probability distributions over functions (28). The algorithm is applied to data deposited in the EM Data Bank (EMDB) (29) and to high-resolution cryo-EM reconstructions of catalase enzymes determined specifically for the purpose of scattering factor estimation. Thanks to recent developments in cryo-EM techniques, the quality of such reconstructions is now high enough to permit the detailed investigation of electron scattering from macromolecules.

## Results

### Scattering model

Under the weak phase approximation, the scattering amplitude of an object *F* ^(*c*)^(**s**) is proportional to the Fourier transform of the scattering potential. For electron scattering, this potential is the ESP *V* (**x**) (1):

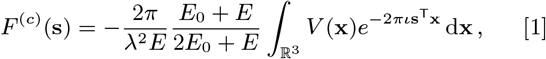

where *E*_0_ is the rest energy of an electron, *E* is the energy of the accelerated electron, and *λ* is the electron wavelength. In order to make the calculation of scattering amplitudes tractable for macromolecules, further assumptions must be made about *V*. One common approximation, which is adopted in this work, is that *V* can be written as a sum of atomic contributions, that is

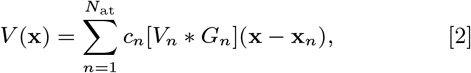

where *V*_*n*_ is the contribution to the ESP of atom *n*, **x**_*n*_ is a 3-vector containing the Cartesian coordinates of the atom, and *c*_*n*_ is the site occupancy. Equation 2 is known as the independent atom model (IAM). The function 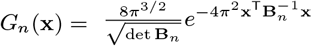 represents the contribution of the ADP of the *n*th atom **B**_*n*_, which models the uncertainty in the atomic position. Here **B**_*n*_ is a positive definite 3 *×* 3 matrix. *V*_*n*_ is further assumed to have spherical symmetry:

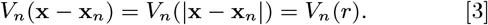

Combining Equations 2 and 3 with the weak phase approximation, *F* ^(*c*)^(**s**) may be written (30)

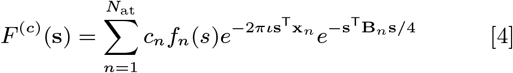

where *s* = |**s**| and *f*_*n*_(*s*) is the atomic scattering factor for atom *n*.

### Atom types

In order to make the scattering factors transferable between different macromolecules, it is assumed that atoms fall into type classes, where all atoms in a type have the same scattering factor. Assuming there are *C* types of atoms, the set of atoms of type *i* is denoted by 𝒜_*i*_ and *f*_*i*_(*s*) is the scattering factor of atoms of this type. Then Equation 4 becomes

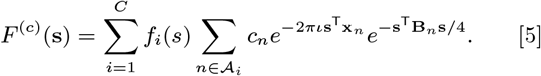

The most widely used form of the IAM is recovered by making each 𝒜_*i*_ correspond to an element, so all atoms of the same element are assumed to scatter in the same way. This model will be referred to as the ‘classical IAM’, and any other scheme for classifying atoms as a ‘generalised IAM’.

Here, the classical IAM is extended by requiring that atoms in the same atom type class are (a) of the same element and further (b) are covalently bonded to the same set of elements. The scheme was further modified to make separate classes for (1) primary amide oxygens in Asn and Gln and (2) carboxyl oxygens in Asp and Glu. Chemical knowledge from the CCP4 Monomer Library (31) and functions available in the package *GEMMI* (32) were used for atom type determination. Throughout this work, a simple notation will be used to refer to atom types, in which bonded neighbours are placed in brackets after the element symbol; for example, C(COO) represents a carbon atom bonded to two oxygen atoms and another carbon atom (Figure 1A).

**Fig. 1.**
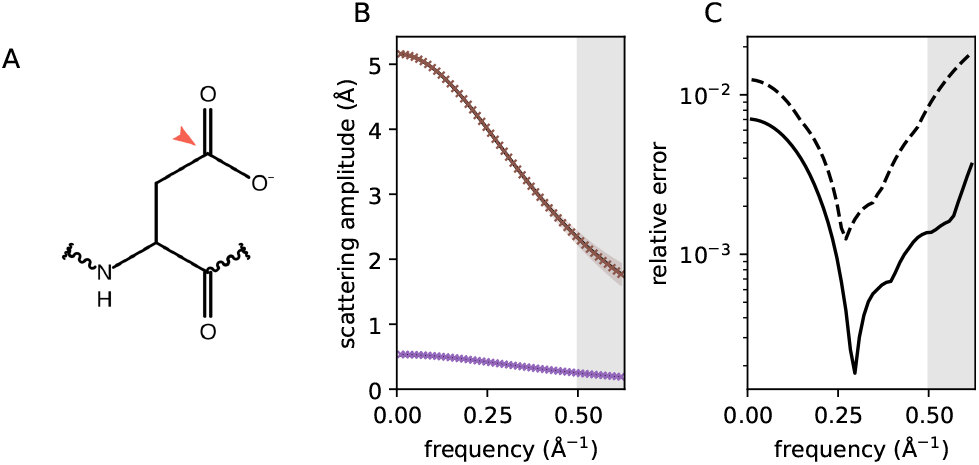
Recovery of tabulated scattering factors from simulated data. (A) Example of classification of atoms into type classes for an Asp side chain. The atom indicated by the red arrow is of type C(COO). (B) The ground truth (solid line) and estimated scattering factors (crosses) for the atom type with the fewest members (S(CC), brown) and the atom type with the most members (H(C), purple) at a simulated resolution of 2 Å. 95% confidence intervals are shown as shaded regions. The region shaded in grey indicates frequencies above the 2 Å resolution cutoff. (C) Mean (solid line) and maximum (dashed line) relative errors as a function of frequency.

### The generalised IAM as a basis set expansion

In a suitable discretised form, *F* ^(*c*)^ in Equation 5 is an expansion over a set of basis functions, in which the coefficients of the expansion are the scattering factors. To show this, first define a ‘frequency bin’ as

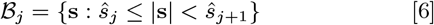

where ŝ_*j*_ are appropriately defined frequency boundaries. A discrete approximation of a scattering factor satisfies

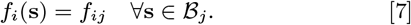

*f*_*ij*_ will become the coefficients of the basis set expansion. The discretised scattering factors are assumed constant within each frequency bin, an approximation that is valid as long as bins are sufficiently narrow. Now define the basis functions:

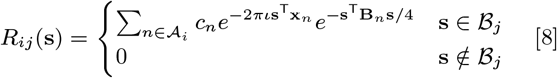

Using Equations 7 and 8, and adopting the Einstein summation convention, Equation 5 may be written

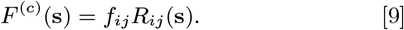

In practice, the observations *F* ^(*o*)^ are located at a set of frequencies (**s**_1_, **s**_2_, …, **s**_*K*_) on a Cartesian grid. The observations may be placed in a vector with elements indexed by *k* such that 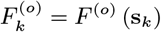. The corresponding scattering amplitudes under the IAM may be obtained from Equation 9 and written as 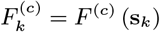. Similarly, the values of the basis on a Cartesian grid may be denoted by *R*_*ijk*_ = *R*_*ij*_ (**s**_*k*_). Then, in the case of observations at discrete frequencies, Equation 9 becomes

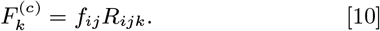

In this work, it is assumed that the parameters of the basis set (atomic coordinates and ADPs) are known prior to inference. They are determined by performing standard atomic model refinement.

### Probabilistic model

The aim of the present work is to estimate a set of scattering factors *f*_*ij*_ from observations, more specifically from the Fourier transform of a cryo-EM map 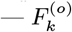. Such scattering factors will be called ‘empirical’ to emphasise their origin. The problem will be formulated within the framework of GP regression. GPs have found widespread applications in machine learning (see (33) for a comprehensive introduction). A GP is an extension of the multivariate normal distribution to infinite-dimensional spaces of functions. Any finite sample of values from a GP follows a multivariate normal distribution with a mean vector ***µ*** and covariance matrix **C**. GPs allow the application of Bayesian techniques to the problem of inferring functions from noisy observations. GPs are also applicable to the case in which the observations are a linear transformation of a latent process (34), a property which will be exploited below.

Assume that each atomic scattering factor in a molecule is drawn from a GP. The domain of the process is frequency, and the GP is real-valued, since atoms are assumed to be radially symmetric. The observations are generated from atomic scattering factors according to Equation 10 with addition of normally distributed observation noise. Formally, define the likelihood as

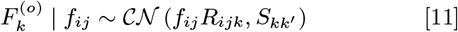

𝒞 𝒩denotes a complex normal distribution with iid real and imaginary parts and 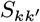 is diagonal. Now place a prior distribution on the scattering factors *f*_*ij*_,

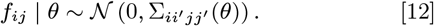

The prior covariance Σ depends on some hyperparameters *θ*. The algorithm does not depend on a particular form of Σ. The covariance function chosen for this study and the procedure used for hyperparameter estimation are described in the SI Appendix.

Since the likelihood and prior are normal, the posterior will also be normal and can be derived by making use of standard results (33, 35). Defining 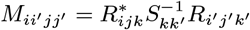and 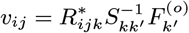

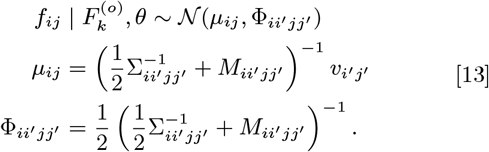

The elements of 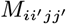 have a simple interpretation as the (noise-weighted) overlap between basis functions *R*_*ij*_ and 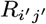. Similarly, the vector *v*_*ij*_ may be interpreted as describing the overlap between a basis function *R*_*ij*_ and the observations *F* ^(*o*)^. The algorithm may be further extended to simultaneously perform inference from multiple cryoEM maps and to make inference more robust to model misspecification (see SI Appendix).

Once *µ*_*ij*_ and the covariance hyperparameters have been estimated, standard results for GPs (33, 35) are used to predict the values of scattering factors at any frequency. The equation to predict the scattering factor at some new frequency 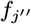 from values *f*_*j*_ available in the training set is

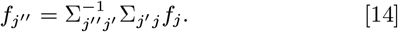

Here, *j* and *j*^*′*^ index frequency bins in the training set, and *j*^*′′*^ indexes frequencies at which the function values will be predicted.

### Performance on simulated data

The algorithm was first applied to the problem of recovering tabulated atomic scattering factors from simulated data. ESP maps were generated from eight sets of atomic coordinates as described in the SI Appendix. Noise was added to the maps to simulate an effective resolution of 2 Å. It was found that scattering factors could be accurately recovered by the algorithm. The recovery of the ground truth is illustrated in Figure 1B, which shows results for the most common (H(C), 84 758 occurrences) and least common (S(CC), 271 occurrences) atom types in the simulated dataset. Figure 1C shows the maximum and mean relative errors in each frequency bin, weighted by the number of atoms in each type class. When considering frequencies below the 2 Å cutoff, the worst case error was 1.2%, while the mean relative error was never above 0.7%.

### Agreement with experimental cryo-EM data

Two training sets of cryo-EM maps and corresponding atomic models were prepared: one containing three high-resolution reconstructions of catalase enzymes (see Materials and Methods), and a second dataset containing 52 entries from the EMDB (prepared as described in the SI Appendix). The inference algorithm was run on the training sets to calculate atomic scattering factors (Figure 2A). Scattering factors from the two training sets were evaluated on a test set consisting of 12 entries from the EMDB (listed in the SI Appendix). Equation 10 was used together with scattering factors calculated from a training set to calculate the molecular scattering amplitude for each test structure. As a baseline, the scattering amplitude was also evaluated using atomic scattering factors from the International Tables (6). The fraction of variance unexplained (FVU, Equation 17) was then used to assess agreement of each model with the observed cryo-EM map. The FVU is the proportion of the data variance that is not explained by the atomic model.

**Fig. 2.**
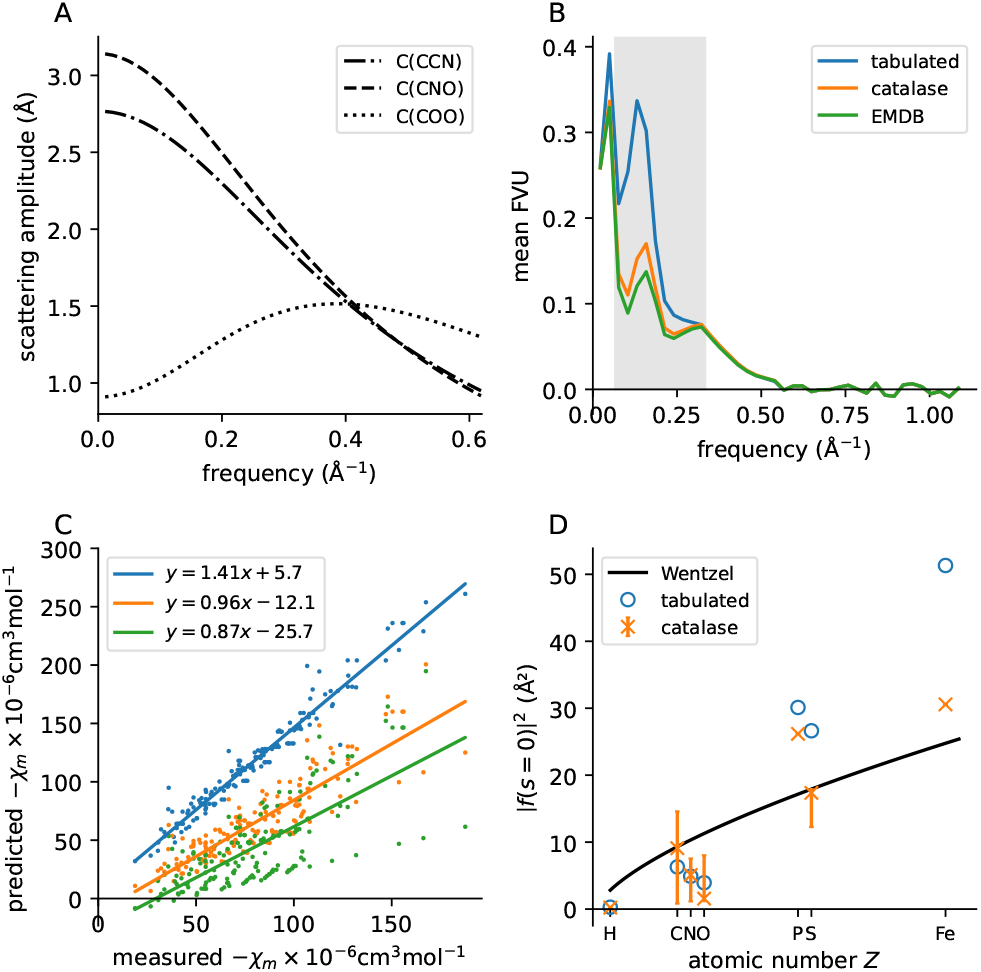
Consistency of empirical scattering factors with experimental data. (A) Scattering factors for three types of carbon atom, calculated from the catalase training set. (B) Mean FVU for structures in the test set, for scattering amplitudes calculated using different sets of scattering factors. The shaded region indicates the frequency range 1/15–1/3 Å ^-1^. (C) Magnetic susceptibility predictions with different sets of scattering factors plotted against measured values. The colour scheme is as in panel B. (D) Differential scattering cross sections at *s* = 0 as a function of atomic number. Predictions from the Wentzel model, tabulated values and values determined from the catalase training set are shown. For catalase, the weighted mean for each element (weighted by the number of examples of each atom type in the training set) is shown as a cross, while the bar gives the range of values.

Figure 2B shows the mean FVU on the test set as a function of frequency. For scattering factors determined from the catalase training set, there is a substantial decrease in the FVU compared to the corresponding metric calculated from tabulated scattering factors. The decrease occurs at low frequency, with the greatest improvement in the frequency range 1/15–1/3 Å^-1^. Scattering factors calculated from the EMDB training set behave similarly, but show a slightly smaller FVU compared to scattering factors from the catalase training set.

### Magnetic susceptibility predictions

In order to further validate the empirical scattering factors, their consistency with measured properties of materials was examined. The molar diamagnetic susceptibility *χ*_*m*_ is a proportionality constant that links the applied magnetic field **H** and the magnetisation **M** as **M** = *χ*_*m*_**H** (36). Under a semiclassical approximation, the susceptibility can be calculated from the atomic scattering factors as (37)

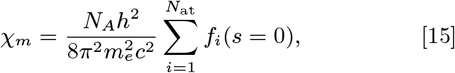

where *N*_*A*_ is Avogadro’s constant, *h* is Planck’s constant, *m*_*e*_ is the electron mass and *c* is the speed of light. The diamagnetic susceptibility is proportional to the mean inner potential (MIP) (37). The MIP is the spatial average of the ESP in a material, and is known to be highly sensitive to the distribution of bonding electrons (38, 39).

Equation 15 was used to calculate *χ*_*m*_ for compounds with measurements available in (36). Empirical scattering factors were first extrapolated to *s* = 0 using Equation 14. Atom typing was performed according to the scheme described above, except that no distinction was made between ***sp*^2^**-hybridised oxygen atoms bonded to carbon in different functional groups (the scattering factor for oxygen on the peptide backbone was used in all cases). The calculation was only performed if all atom types present in a molecule were also present in the catalase training set. Data from a total of 177 compounds were used in the analysis.

Figure 2C shows the relationship between predicted and measured values of *χ*_*m*_ using tabulated scattering factors, as well as empirical scattering factors from the two training sets. *χ*_*m*_ is systematically overestimated by the tabulated scattering factors, a phenomenon which has been reported previously for estimates of the MIP (40–43). Linear least-squares regression of empirical against calculated values was used to measure this systematic error. The error is reflected in the slope fitted by linear regression, 1.41, and root mean square deviation (rmsd) value of 41.5 *×* 10^−6^cm^3^mol^−1^. On the other hand, when *χ*_*m*_ is estimated using empirical scattering factors from the catalase training set, the equation of the line is *y* = 0.96*x* − 12.1 and the rmsd is 22.4 *×* 10^−6^cm^3^mol^−1^. Using empirical scattering factors from the catalase training set therefore decreases the error in estimation of *χ*_*m*_ compared to the tabulated scattering factors. However, it results in systematic underestimation of this quantity.

Finally, using scattering factors calculated from the EMDB training set did not result in an improvement over using tabulated scattering factors (equation: *y* = 0.87*x* − 25.7, rmsd = 40.5 *×* 10^−6^cm^3^mol^−1^). A possible explanation is the presence of poorly resolved regions in maps in the EMDB. A decrease in the signal power at some atoms in an atom type class would result in a decrease in its estimated scattering amplitude, in turn causing underestimation of *χ*_*m*_. Other factors that may affect the results include radiation damage to the specimen, errors in the deposited atomic model, and errors associated with extrapolation of scattering factors to *s* = 0. The latter in particular may explain the large spread in the predicted *χ*_*m*_ values. In light of this analysis, only scattering factors from the catalase dataset were used for the atomic model refinement outlined in the following section.

The dependence of the zero-frequency scattering on the atomic number *Z* was also investigated and compared to the Wentzel atom model (44), a simple description of the screening action of electrons on the nuclear ESP. The Wentzel model predicts that the differential scattering cross section (defined as the square of the scattering factor) at *s* = 0 is proportional to *Z*^2*/*3^ (1, Equation 5.34). It was found that the dependence of the cross section on atomic number is qualitatively similar for tabulated scattering factors and those determined from the calatalase training set (Figure 2D): the cross section decreases in the progression C *>* N *>* O, in contrast to the Wentzel model, which predicts an increase. The decrease replicates the trend in the covalent radii of these three elements (45).

### Atomic model refinement with empirical scattering factors

A major application of scattering models in cryo-EM is in atomic model refinement (47). Atomic model refinement with a generalised IAM was implemented in the program *Servalcat* (48). The 12 structures in the test set were then refined using empirical scattering factors from the catalase training set. The results were compared against refinements with the scattering model currently used by *Servalcat*, which converts the electron density calculated using a classical IAM into an electrostatic potential by means of the Mott-Bethe formula (49, 50).

The atomic coordinates obtained using empirical scattering factors do not differ substantially from those produced by a standard model refinement, with a maximum structure rmsd of 0.18 Å. Across the whole test set, only 12 residues contained atoms with positions differing by more than 1 Å from the baseline. Following refinement, *F* ^(*o*)^ − *DF* ^(*c*)^ difference maps were computed for each structure in the test set. The most notable effect of using empirical scattering factors compared to the current model is a decrease in the size of negative difference density on Asp/Glu side chains compared to the baseline (Figure 3A-D).

**Fig. 3.**
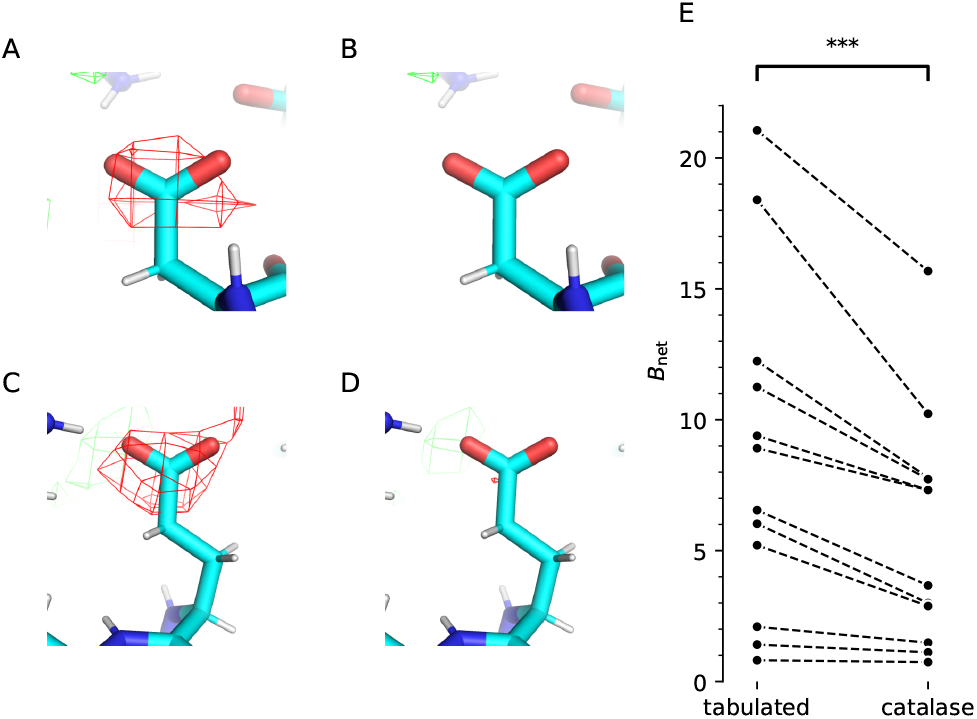
Atomic model refinement and *F* ^(*o*)^ − *DF* ^(*c*)^ difference map calculation using empirical scattering factors. (A-D) Comparison of difference maps calculated with tabulated (A, C) and empirical (B, D) scattering factors. Maps are contoured at the 4*σ* level. (A, B) show Asp46 and (C, D) show Glu47 from deposition EMD-13937 (connexin 26). Panels A-D were prepared using *PyMOL* (46). (E) *B*_net_ for the 12 structures in the test set after refinement using tabulated scattering factors (left column) and empirical scattering factors from the catalase training set (right column). *RABDAM* was used for calculating *B*_net_. Details of the test structures are given in the SI Appendix.

In contrast to the positions of atoms, which remain almost unchanged, ADPs decrease in generalised IAM refinement compared to the baseline. Since the absolute scale of ADPs is arbitrary (51), the following procedure was adopted for analysing ADP differences: the ADP difference was calculated for each atom in the structure, a *z*-score was calculated for each atom according to this distribution, and atoms with *z <* −3 were analysed. Most of these atoms (94.9%) are from Asp/Glu carboxyl groups or the sulphur atom in Cys. This trend was further investigated by calculating the *B*_net_ metric, which measures the magnitude of ADPs of Asp/Glu atoms compared to atoms in similar packing environments (52). The software package *RABDAM* calculates this metric (53). *B*_net_ decreased for all structures in the test set, and the decrease was significant (Figure 3E; *t* = −4.27, *p* = 0.0007). The analysis demonstrates that empirical scattering factors absorb common features of cryo-EM maps, in particular the poor visibility of carboxyl groups. However, as discussed below, it should be noted that all cryo-EM data suffers from the effects of beam-induced damage to the sample, a factor that must be taken into account when interpreting the results.

## Discussion

This article describes the development and application of a method for the inference of atomic scattering factors from cryo-EM maps. The inference algorithm presented above is based on reformulating the IAM, which is the most commonly-used forward model for the ESP in cryo-EM data analysis, as an expansion over a fixed basis. The inverse problem, that is inference of scattering factors from an ESP distribution reconstructed by cryo-EM SPA, can then be solved within the probabilistic framework of GP regression. Furthermore, the ‘classical’ IAM may be viewed as a special case of any forward model that expresses the ESP as a sum of atomic contributions. This broad class of models is referred to as the ‘generalised’ IAM. By carefully choosing the assignment of atoms to type classes in the generalised IAM, the scattering factors inferred by the algorithm can be made to reflect charge redistribution due to covalent bonding. It should be noted that the algorithm does not depend on a particular atom typing scheme. For example, atom types defined in various molecular mechanics force fields could be used instead of the automatic procedure described above. Whatever atom typing scheme is used, it must balance the requirement for classes that are fine enough to reflect the local chemical environment, but at the same time contain enough examples to allow accurate estimation of the model parameters.

Scattering factors determined by the inference algorithm show improved agreement with cryo-EM data and measured magnetic susceptibility values. The latter observation suggests that scattering factors derived from high-resolution cryo-EM reconstructions may find application in calculation of the MIP, a quantity important in electron holography and other EM imaging modes (54). However, the accuracy of scattering factors near *s* = 0 is currently limited due to reasons already discussed. The new scattering factors are also compatible with existing methods for atomic model refinement. While usage of empirical scattering factors does not lead to significant changes in the positions of atoms, it does affect ADPs. Using scattering factors that take into account bonding effects may lead to more accurate determination of ADPs from cryo-EM data, unlocking further information about atomic motion and disorder in biological macromolecules. Reference-based template matching (55) is another application that may benefit from improved electron scattering models. Template matching locates a macromolecule in micrographs by searching for regions of the image that correlate with a precomputed map of the macromolecular ESP. A more accurate scattering model would increase the correlation between the precomputed ESP and the observations, improving detectability.

Although there are substantial improvements in the agreement between the cryo-EM 3D maps and the models in the frequency range 1/15–1/3 Å^-1^, there are features in the individual atom plots indicating scope for further improvement. These are mainly due to the limitations of the experimental data. There are at least three potential improvements. One of these is that the hydrogen atom coordinates have not been individually refined to fit into the densities, but have been extrapolated from the non-hydrogen positions; higher-resolution maps would be desirable in order to obtain accurate rotational angles for terminal methyl groups such as those in valine and leucine, and for hydrogen-bonded atoms. As a result the hydrogen scattering factors may be underestimated. Also the trend in low-frequency scattering of C *>* N *>* O (in contrast to the trend in atomic number of C *<* N *<* O) is now satisfyingly less than it is for the tabulated IAM values, but it may still be influenced by negative charges on some oxygens, positive charges on some nitrogens, more accurate coordinates for the carbons, as well as increased flexibility and increased radiation damage to some side-chain atoms, especially Glu. There is therefore a need for a higher-resolution structure with much less radiation damage. A larger experimental set of cryo-EM images (possibly ten times larger than a typical dataset) would allow the structure to be obtained at lower electron doses, before damage is manifested. Concurrently, the accuracy of scattering factors derived from publicly-available cryo-EM datasets could be improved by careful rebuilding of the corresponding atomic models.

The scattering amplitude is a quantity that occurs naturally in physical theories of scattering, but is difficult to interpret intuitively. In particular, qualitative descriptions of biochemical processes are often based on some notion of ‘charge’ or ‘polarity’. The charge of an atom in a molecule is not a physical observable, but various methods exist to derive numerical values from the molecular electron density or ESP (reviewed in (56)). Since the empirical scattering factors in this work are also derived from the ESP, it should be possible to explain their behaviour with reference to atomic partial charges. However, the task is not trivial: the empirical scattering factors in this work are well-defined at *s* = 0, therefore correspond to atoms that are formally neutral. A possible avenue for defining partial charges based on empirical scattering factors is to convert them to atomic ESP distributions, then compare the ESP to that of an isolated neutral atom.

A number of authors have also claimed that it is possible to assign partial charges to atoms based on their appearance in cryo-EM maps, focusing in particular on carboxyl groups, which are often not visible in reconstructions of the ESP (20, 23, 24). On the other hand, carboxyl groups are known to be radiation-sensitive, an effect that has been well-documented in X-ray diffraction (XRD) experiments (57, 58). Either of these phenomena (charge or radiation damage) could feasibly explain the poor visibility of carboxyl groups in cryo-EM maps. The present work shows that the scattering factor for carbon in a carboxyl group differs at low frequency from that of carbon atoms on the main chain (Figure 2A). However, it is unclear at this stage whether the difference is due to charge or radiation damage. The scattering properties of carboxyl groups may be further investigated by *ab initio* calculation of the ESP of small molecules, then estimation of scattering factors from the calculated ESPs. The resulting scattering factors should show the effects of charge, but not radiation damage. Alternatively, modelling of radiation damage may be done jointly with inference of scattering factors, either as a preliminary ‘zero-dose extrapolation’ step (59) or within a more complex model of scattering that accounts for changes in specimen structure with increasing electron dose.

Finally, some features of the macromolecular ESP have not been addressed in this work. Firstly, no attempt has been made to model the solvent contribution to the ESP. In cryo-EM, the macromolecule is suspended in vitreous ice, which is a dielectric material and therefore is polarised by the charge distribution of the solute (60). The induced charge contributes to the ESP. The contribution of the solvent is expected to be most significant at low frequency (61). Secondly, the assumption that atoms are spherical is problematic in the case of hydrogen, whose electron density is polarised along the *X* —H covalent bond. It is known from XRD studies that more sophisticated models of hydrogen atoms lead to more accurate values for the positions and ADPs of these atoms compared to the IAM (62–64). With the increasing availability of high-resolution reconstructions, cases in which accurate models of hydrogen atoms are required will become more common in cryo-EM. Aspherical hydrogens could be incorporated into the GP regression algorithm by a suitable modification of the forward model, for example by introduction of an operator that deforms an underlying spherical scattering factor.

## Materials and Methods

### Catalase structure determination

High-resolution electrostatic potential maps were determined by cryo-EM single-particle reconstruction (65) for catalase from three species: *Rhizobium radiobacter* (formerly *Agrobacterium tumefaciens*), *Micrococcus luteus* (formerly *Micrococcus lysodeikticus*), and *Homo sapiens* (erythrocyte cells). Catalases from *R. radiobacter* and *M. luteus* were chosen following a review of publications to search for stable protein targets for cryo-EM that included as many atom types as possible, for example Fe and P present in cofactors. The selection of protein targets will be discussed further in (66).

Full details of the structure determination will be given in (66). In brief, thin films of catalase in solution were vitrified on HexAuFoil supports (59, 67) (produced in house and purchased, Quantifoil). The grids were treated with a 9:1 Ar:O_2_ plasma to render the surface hydrophilic (60-90s, 30 sccm, 40W forward power) (Fischione, 1070 Nanoclean). Grids were plunged using a manual plunger of the Talmon type (68) in a 4°C cold room, equilibrated to greater than 90% relative humidity. For each specimen, a 3 uL droplet of 15-20 mg/mL protein with 5 mM CHAPSO detergent was applied to the foil side of a plasma-treated grid and blotted with filter paper (Whatman No. 1) from the foil side for 15-30 seconds. The grid was then immediately plunged into liquid ethane held at 93K in a temperature controlled cryostat (69).

Movies were acquired on a Titan Krios microscope operating at 300 kV using a Falcon 4i detector in counting mode. The nominal magnification was 155 kX, with a pixel size of 0.512 Å/pixel calibrated using the Au reflections present in a selection of acquired images due to the presence of support foil within the field of view (70). A 100 *µ*m objective aperture was used during data collection. Movies were acquired with 3.0 s exposure time, and the flux was set to 18 eÅ^−2^s_−1_ giving a total fluence of 54 eÅ^−2^.

The electrostatic potential maps were reconstructed from the acquired movies using *RELION-4* (71). First the movies were motion corrected with *MotionCor2* (72), and the CTFs were fitted using *CTFFIND4* (73). Particles were manually picked and used to train a *Topaz* (74) model which was then used for autopicking. *RELION-4* was used for 2D and 3D classification, 3D refinement, CTF refinement, particle polishing, and post-processing.

Models were built following rigid-body docking of PDB depositions into the sharpened maps from *RELION*: 1GWE expanded to the tetramer into the maps for *R. radiobacter* and *M. luteus* catalases, and 7P8W into the map for the human catalase. The models were then manually adjusted, with sequence correction for *R. radiobacter*. Docking and rebuilding were performed in *Coot* (75). The models were refined against the unsharpened halfmaps using *Servalcat*.

### Test metrics

Performance of the empirical scattering factors on the test set was evaluated using the fraction of variance unexplained (FVU), given by

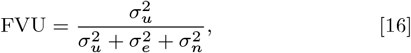

where 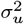 is signal variance unexplained by the model, 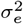 is signal variance explained by the model, and 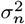 is the noise variance. It is assumed that the explained and unexplained signal are uncorrelated. It can be shown that the FVU is related to the Fourier shell correlation (FSC) as

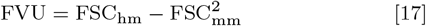

where FSC_hm_ is the FSC between two halfmaps and FSC_mm_ is the FSC between the model and one of the halfmaps.

### Atomic model refinement

Refinement with a generalised IAM was implemented in *Servalcat*. Most atomic model refinement programs in macromolecular crystallography and cryo-EM (including *Servalcat*) use a sum-of-Gaussians parametrisation of the atomic scattering amplitudes (6, 30):

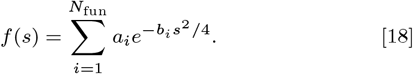

The parametrisation above with *N*_fun_ = 5 was fitted to the empirical scattering factors by minimising a least-squares objective with L2 regularisation applied to the parameters *a*_*i*_. The coefficients *b*_*i*_ were constrained to be positive. Minimisation was performed using L-BFGS (76).

Structures in the test set were re-refined for 10 refinement cycles using the empirical scattering factors and using tabulated scattering factors. Tabulated scattering factors were used for atom types which were not present in the training set.

## Data, Materials and Software Availability

The ESP maps of catalase are available in the EMDB under entries EMD-55403, EMD-55404 and EMD-55405. The atomic models are deposited in the Protein Data Bank under entries 9T0K, 9T0L and 9T0M. Parameters of scattering factors and the simulated dataset are available on Zenodo (https://zenodo.org/records/17084047) (77). The code for scattering factor estimation is available on Github (https://github.com/as2875/sffit).

## ACKNOWLEDGMENTS

We thank J. Grimmett, T. Darling and I. Clayson for maintaining the LMB Scientific Computing facility; we thank the LMB EM facility staff for access to equipment. A.S. thanks S.H.W. Scheres for advice on algorithm design. This work was supported by the Medical Research Council under Grant Nos. MC UP A025 1012 (G.N.M.) and MC UP 120117 (C.J.R.). H.W. is supported by an Astex Pharmaceuticals Sustaining Innovation Postdoctoral Fellowship.

