## Supporting Information Appendix for "Measurement of atomic scattering factors by cryo-electron microscopy"

#### This PDF file includes:

Supporting text
Figs. S1 to S2
Table S1
SI References

### Supporting Information Text

**Form of the covariance function.** In this work, two assumptions are made to construct a tractable covariance $\Sigma_{ii'jj'}$. Firstly, it is assumed that the covariance factorises into 'frequency' and 'atom type' terms, and scattering factors for different atom types are independent, that is $\Sigma_{ii'jj'} = \delta_{ii'}\Sigma_{jj'}$. The second assumption is that a scattering factor $f(s)$ may be expressed as a non-stationary convolution of a Gaussian white noise process $\phi(b)$,

$$f(s) = \int_0^\infty \phi(b) e^{-bs^2} u(b)\,\mathrm{d}b\,, \tag{S1}$$

where $u(b)$ is the scale of the noise process. Then

$$\begin{aligned} \mathbb{E}\left[f(s)\right] &= 0 \\ \mathrm{cov}\left[f(s), f(s')\right] &= \int_0^\infty \int_0^\infty \mathbb{E}\left[\phi(b)\phi(b')\right] e^{-bs^2} e^{-b's'^2} u(b)u(b')\,\mathrm{d}b\,\mathrm{d}b' \\ &= \int_0^\infty e^{-bs^2} e^{-bs'^2} u^2(b)\,\mathrm{d}b \end{aligned} \tag{S2}$$

Further supposing that $u^2(b)/K = \frac{1}{\beta}e^{-b/\beta}$ is the PDF of an exponential distribution, where $K$ is a normalising constant, the covariance becomes

$$\mathrm{cov}\left[f(s), f(s')\right] = K\left(1 + \beta(s^2 + s'^2)\right)^{-1}. \tag{S3}$$

This construction may be justified as a continuous extension of a discrete summation of Gaussians, the latter being the standard parametrisation for X-ray and electron scattering factors used in macromolecular crystallography (1, 2). For the discretised observation model, $\Sigma_{jj'} = \mathrm{cov}\left[f(s_j), f(s_{j'})\right]$, where $s_j = \frac{d_j + d_{j+1}}{2}$.

**Hyperparameter estimation.** The noise variance $S_{kk'}$ is estimated in frequency bins using maximum likelihood (ML), as in *REFMAC* (3) and *Servalcat* (4). The ML estimator has a closed form. The noise variance $\sigma_j$ in $\mathcal{B}_j$ is

$$\sigma_j = \frac{1}{N_j} \sum_{k:\mathbf{s}_k \in \mathcal{B}_j} \left| F_k^{(o)} - F_k^{(c)} \right|^2, \tag{S4}$$

where $N_j$ is the number of coefficients in $\mathcal{B}_j$ and $F_k^{(c)}$ is calculated using tabulated scattering factors. Then

$$S_{kk'} = \delta_{kk'} \sum_{j=1}^{N_{\mathrm{bin}}} 1_{\mathcal{B}_j}(\mathbf{s}_k)\sigma_j, \tag{S5}$$

where $1_{\mathcal{B}_j}$ is the indicator function of $\mathcal{B}_j$ and $N_{\mathrm{bin}}$ is the number of frequency bins.

The parameters of the covariance $\theta = \{K, \beta\}$ are estimated by maximising the marginal likelihood of the observations given the parameters, where marginalisation is over values of the scattering factors. This method is an application of the evidence approximation (5). For the model adopted in this work, the negative marginal log-likelihood is

$$\begin{aligned} -\log p(F_k^{(o)} \mid \theta) = {} & F_k^{(o)} S_{kk'}^{-1} F_{k'}^{(o)} - v_{ij}\left(\frac{1}{2}\Sigma_{ii'jj'}^{-1} + M_{ii'jj'}\right) v_{i'j'} \\ & + \frac{1}{2}\log\det\left(\frac{1}{2}\Sigma_{ii'jj'}^{-1} + M_{ii'jj'}\right) + \frac{1}{2}\log\det\left(\Sigma_{ii'jj'}\right) + \log\det\left(S_{kk'}\right) + \text{const.} \end{aligned} \tag{S6}$$

The inference algorithm first finds the hyperparameters $\theta$ that minimise the negative marginal log-likelihood in Equation S6 using the iterative optimiser L-BFGS (6). It then computes the posterior expectation $\mu_{ij}$ from Equation 13, which is the solution to a linear system. In order to speed up the hyperparameter estimation, the quantities $M_{ii'jj'}$ and $v_{ij}$ are precomputed.

**Inference from multiple cryo-EM maps.** The probabilistic model can be extended to datasets containing multiple cryo-EM maps. Maps are assumed to be conditionally independent given the corresponding atomic models. Indexing the map by $l$, the prior remains unchanged while the likelihood becomes

$$F_{kl}^{(o)} \mid f_{ij} \sim \mathcal{CN}\left(f_{ij} R_{ijkl}, S_{kk'll'}\right), \tag{S7}$$

where $S_{kk'll'}$ is diagonal. The latter property follows from the conditional independence assumption. On redefining $M_{ii'jj'} = R_{ijkl}^* S_{kk'll'}^{-1} R_{i'j'k'l'}$ and $v_{ij} = R_{ijk}^* S_{kk'll'}^{-1} F_{k'l'}^{(o)}$, Equation 13 still holds.

To ensure consistency between datasets, $F^{(o)}$ is scaled to $F^{(c)}$ isotropically in frequency bins. The scaling parameter is estimated by ML, as in *REFMAC* and *Servalcat*. The ML estimator for the multiplicative factor $D_j$ that scales $F^{(c)}$ onto $F^{(o)}$ is

$$D_j = \frac{\sum_{k:\mathbf{s}_k \in \mathcal{B}_j} F_k^{(o)} F_k^{(c)*}}{\sum_{k:\mathbf{s}_k \in \mathcal{B}_j} \left| F_k^{(c)} \right|^2}, \tag{S8}$$

where $F_k^{(c)}$ is again calculated using tabulated scattering factors.

**Robust inference.** The scattering model adopted in this work does not precisely describe the data-generating process, since it makes a number of simplifying assumptions (sphericity of atoms, absence of a solvent contribution, and so on). The model is therefore misspecified. Inference is made more robust to this misspecification by using a 'power likelihood' (7), which is constructed by multiplying $M_{ii'jj'}$ and $v_{ij}$ by a constant $\alpha < 1$. The value of $\alpha$ is estimated by a validation set method. For a given value of $\alpha$, the hyperparameters and scattering factors are estimated using only coefficients with frequencies above 1/5 Å$^{-1}$. The estimated scattering factors are then used to compute values of $F^{(c)}$ for the held-out frequencies. The value of $\alpha$ maximising the correlation between the calculated and observed coefficients is found by performing a line search.

The strength of signal due to electron scattering varies across a cryo-EM map (8). In any given map, some atoms will therefore be poorly resolved. In order to reduce the influence of such atoms on the inference algorithm, they are not used for estimation of the scattering factors. Atoms with an ADP greater than 1.5 times the interquartile range above the median ADP of atoms in the same structure are considered outliers. The threshold translated to exclusion of 12.9% of atoms in the catalase dataset and 13.4% of atoms in the EMDB dataset. For the catalase dataset, the ADP cutoff values were 92.6 Å$^2$ for human erythrocyte catalase, 77.9 Å$^2$ (*Micrococcus luteus*) and 86.9 Å$^2$ (*Rhizobium radiobacter*). The scattering contribution of excluded atoms is calculated using tabulated scattering factors and subtracted from the map before estimation.

**Preparation of datasets.** Simulated maps were generated from eight experimental structures of catalase enzymes (PDB depositions 8EL9, 8PVD, 8SGV, 8WZH, 8WZJ, 8WZK, 8WZM, 9BDJ). Ligands, waters and hydrogen atoms were removed and isotropic ADPs were randomised before simulation by drawing their values from an inverse-gamma distribution (9). *GEMMI* and tabulated scattering factors (1) were used to generate ESP maps. Gaussian noise was added to the simulated maps in Fourier space. The power spectrum of the noise was chosen to (a) ensure that the FSC at 1/2 Å$^{-1}$ was equal to 0.143 and (b) produce sigmoidal FSC curves that are typical of high-resolution cryo-EM reconstructions.

The EMDB dataset is a subset of maps deposited in the EMDB with (a) reported resolution of 2 Å or better, (b) an associated PDB deposition with no missing link records and (c) deposited half-maps. Model refinement was performed for each of the maps in the dataset using *Servalcat*. Maps with an average map-model FSC after refinement of below 0.6 were excluded, these were found to have large unmodelled or flexible regions. The box size of each map was then trimmed to within 20 Å of atoms in the model using *Servalcat*. Maps that had more than $300^3$ voxels after trimming were excluded. The final dataset consisted of 64 maps, which were randomly split into training (52 maps) and test (12 maps) datasets.

Details of the test set are given in Table S1. The following EMDB entries were used in the training set: EMD-60655, EMD-21024, EMD-29357, EMD-42164, EMD-51522, EMD-28259, EMD-15312, EMD-11657, EMD-26756, EMD-31910, EMD-17508, EMD-11638, EMD-45626, EMD-45628, EMD-23749, EMD-45192, EMD-16783, EMD-16788, EMD-17522, EMD-27316, EMD-25201, EMD-28759, EMD-26996, EMD-16785, EMD-11233, EMD-16786, EMD-27285, EMD-41150, EMD-17129, EMD-16787, EMD-26801, EMD-19938, EMD-19881, EMD-17521, EMD-61728, EMD-17510, EMD-16789, EMD-42775, EMD-60391, EMD-28812, EMD-16531, EMD-16784, EMD-42793, EMD-44299, EMD-21951, EMD-51230, EMD-61727, EMD-14332, EMD-27286, EMD-17511, EMD-41149, EMD-26757.

**Table S1. Details of structures in the test set**

| EMDB ID | reported resolution (Å) | description |
|---|---|---|
| EMD-10101 | 1.84 | apoferritin from mouse |
| EMD-13937 | 1.9 | human connexin 26 dodecamer at 90mmHg PCO2, pH7.4 |
| EMD-61729 | 1.83 | ferritin variant R63MeH/R67MeH with Cu(II) |
| EMD-28758 | 2.0 | calcitonin receptor in complex with Gs and pramlintide analogue peptide San45 |
| EMD-51521 | 1.99 | glucose/xylose isomerase from *Streptomyces rubiginosus* with cobalt ions in the active site |
| EMD-14705 | 1.77 | human apoferritin obtained from ssDNA coated grid |
| EMD-29664 | 1.98 | wild-type GAPDH |
| EMD-28269 | 1.68 | mouse apoferritin heavy chain without zinc determined using single-particle cryo-EM with Apollo camera |
| EMD-27833 | 1.98 | helical arch of BIRC6 (from local refinement 1) |
| EMD-60915 | 1.51 | mouse heavy-chain apoferritin |
| EMD-61726 | 1.78 | ferritin variant R63BrThA/E67BrThA |
| EMD-17520 | 1.8 | CAK in complex with inhibitor ICEC0943 |

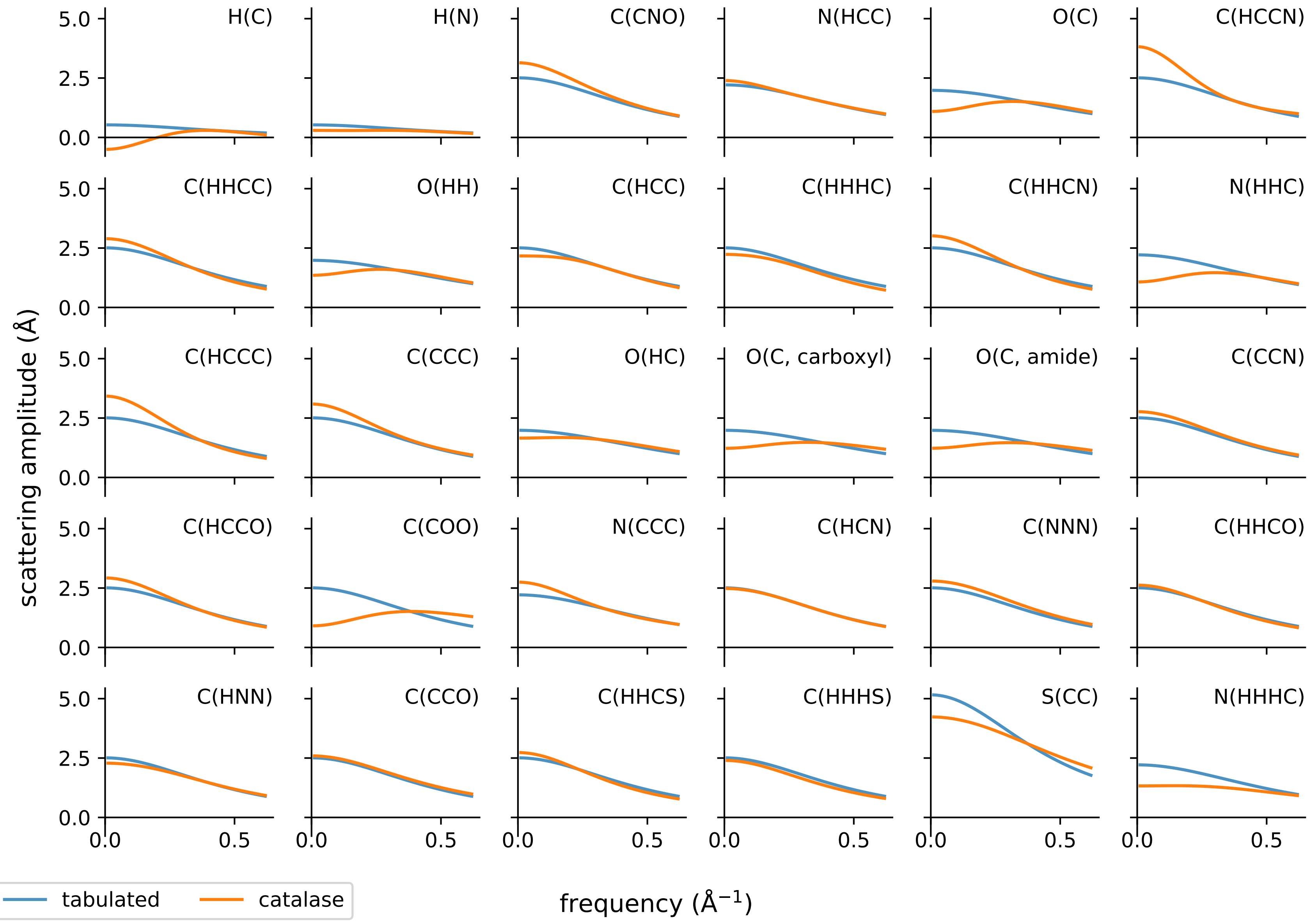


**Fig. S1.** Tabulated scattering factors (blue) and scattering factors determined from the catalase training set (orange) for the 30 most common atom types in the catalase training set.

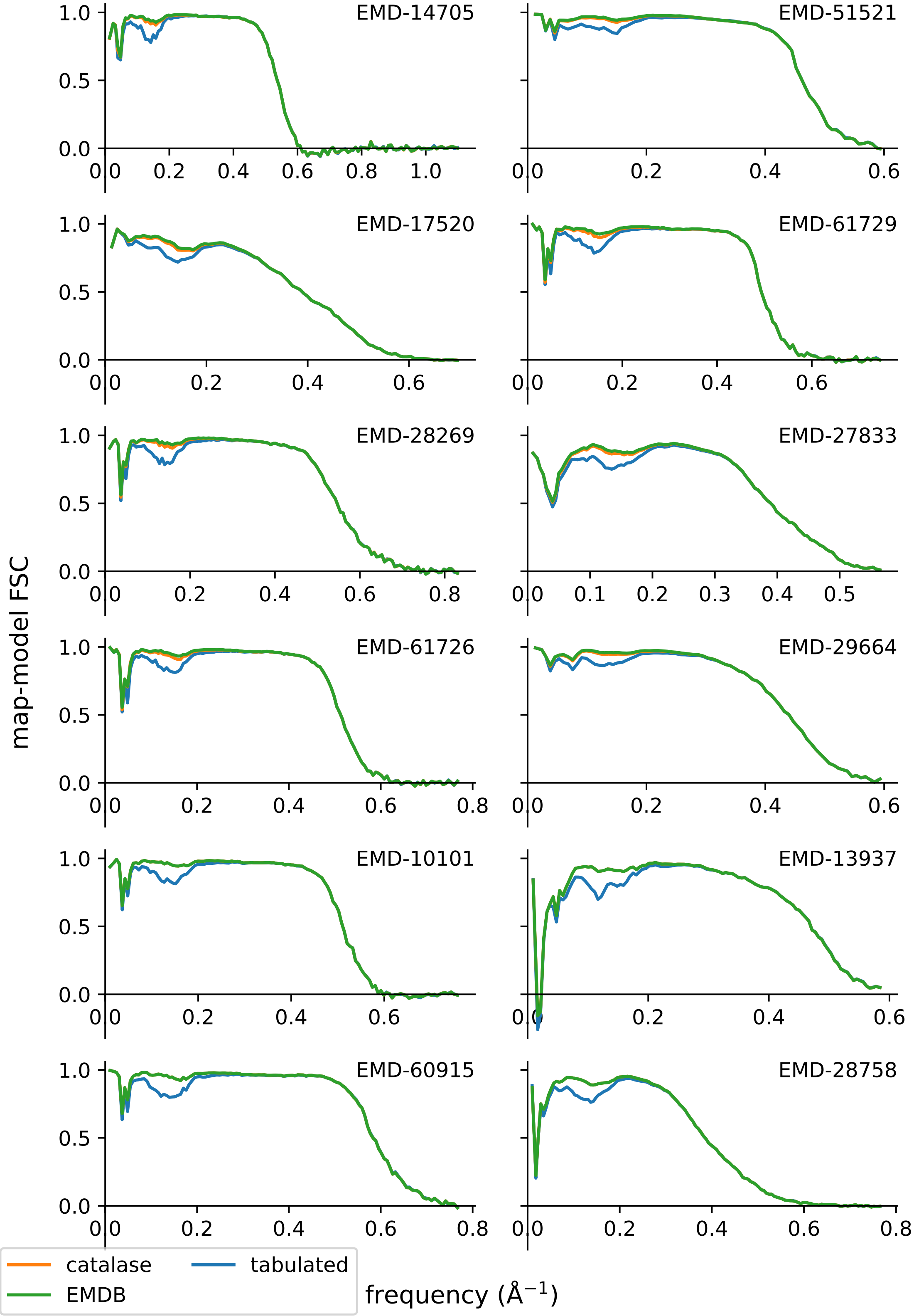


**Fig. S2.** Map-model FSC for structures in the test set, for scattering amplitudes calculated using different sets of scattering factors.
